## Supporting info for "A successional shift enhances stability in ant symbiont communities"

### Supplementary Information: A successional shift enhances stability in ant symbiont communities

### 1 Model selection and diagnostics

Table S1: Summary measures of model selection and model fit. WAIC stands for Widely-Applicable Information Criterion, $R^2$ gives the explanatory power of each model. Predictors: con = connectivity, can = canopy openness, quadratic terms are indicated with $^2$.

| Model | Predictors | WAIC | $R^2$ |
|---|---|---|---|
| 1 | age+con+moisture+moisture$^2$+pH+can | 511.3 | 0.51 |
| 2 | age+con+moisture+pH+pH$^2$+can | 611.6 | 0.47 |
| 3 | age+con+moisture+moisture$^2$+pH+pH$^2$+can | 502.9 | 0.47 |
| 4 | age+con+moisture+moisture$^2$+pH+pH$^2$+can+can$^2$ | 673.5 | 0.49 |
| 5 | age+con+moisture+moisture$^2$+pH+can+can$^2$ | 490.7 | 0.55 |
| 6 | age+con+moisture+pH+pH$^2$+can+can$^2$ | 359.4 | 0.54 |
| 7 | age+con+moisture+pH+can+can$^2$ | 339.3 | 0.55 |
| 8 | age+con+moisture+pH+can | 340.2 | 0.55 |

Model 7 was retained based on the lowest WAIC value. This model also had the highest $R^2$. The MCMC convergence of model 7 was satisfactory. The trace plots displayed an irregular pattern with similar running means across all chains. The potential scale reduction factors were smaller than 1.1 and the effective sample size was close to the actual sample size of 4000 (each of the 4 chains has 1000 samples).

The predictive power of the retained model ($R^2 = 0.10$), was lower than the explanatory power ($R^2 = 0.55$). This relatively low predictive power may be related to a relatively small number of sampled nests ($N = 51$). Since our research places a greater emphasis on inference rather than prediction, we have confidence in the robust support for our conclusions. This is evident by the strong statistical support of relevant parameter estimates, despite the low predictive power of the model.

#### 2 Supplemental figure

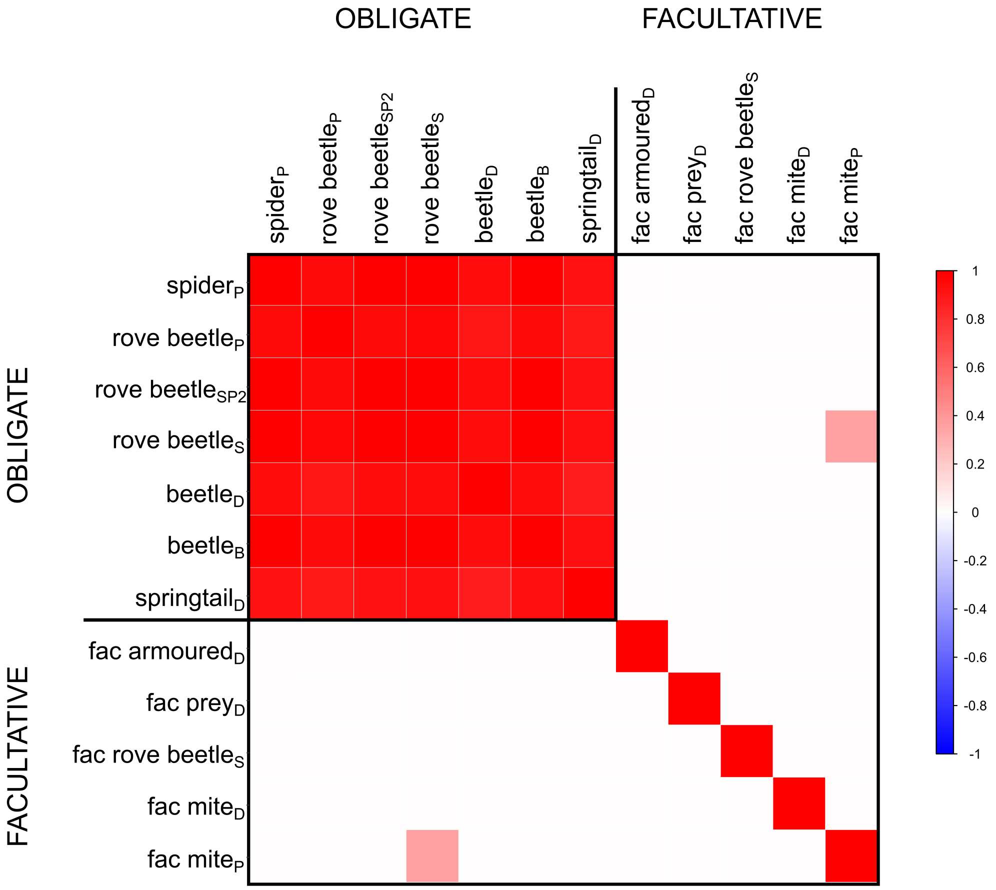


Figure S1: Residual association in abundance shown for all pairs of functional groups after controlling for the predictors in the retained model 7. The functional groups are grouped in obligate and facultative red wood ant associates. Each matrix cell corresponds to a pair of functional groups and the gradient colours indicate the estimated association strength measured at correlation scale. Only associations with at least 95% posterior probability are displayed. Red colors depict positive residual associations among functional groups, no negative residual associations were detected.

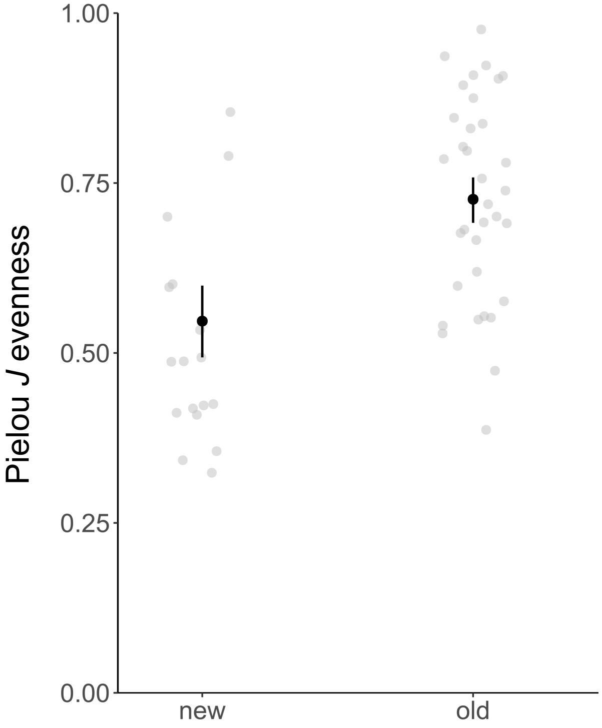


Figure S2: Evenness of symbiont communities in new and old nests as measured by the Pielou's evenness index $J$, beta regression, P $<$ 0.001)
